# Inhibition of Talin-induced Integrin Activation by a Double-hit Stapled Peptide

**DOI:** 10.1101/2022.08.12.503760

**Authors:** Tong Gao, Eun-ah Cho, Pingfeng Zhang, Jinhua Wu

## Abstract

Integrins are ubiquitously expressed cell-adhesion proteins. Talin is required for integrin activation through an inside-out signaling pathway, during which talin is recruited to the plasma membrane (PM) by RAP1 directly or through its effector, RAP1-Interacting Adaptor Molecule (RIAM). RIAM also activates talin from autoinhibition by binding to talin head domain. A helical talin-binding segment (TBS) in RIAM mediates both talin activation and recruitment by binding to two distinct sites in talin head and rod domains respectively. The bi-specificity of the TBS fragment allows us to develop a new strategy to suppress talin-induced integrin activation through a “double-hit” approach. We designed an experimental peptidomimetic inhibitor by engineering a hydrocarbon “staple” in the helical TBS fragment to mask the integrin binding site in the talin head. The stapled peptide (S-TBS) exhibits a stronger binding affinity with talin and inhibits talin:integrin interaction. Crystal structure of S-TBS in complex with talin rod domains reveals an interface that overlaps with the TBS-binding site in talin. Consistently, S-TBS also exhibits an inhibitory effect on TBS:talin-rod interaction during the PM recruitment of talin. Importantly, S-TBS possesses excellent cell permeability and inhibits integrin activation in a talin-dependent manner. Hence, our results present a novel approach to design a new class of intracellular inhibitors targeting integrin functions.

## Introduction

Integrins are heterodimeric adhesion receptors that transduce bi-directional signals across the plasma membrane (PM). A cytoplasmic signaling cascade known as inside-out signaling pathway triggered by growth factor stimulation induces conformational changes in the integrin ectodomains, leading to integrin activation (1). Activated integrins can bind extracellular ligands and trigger downstream signaling that induces cell adhesion, spreading, migration, proliferation, and survival (2). The roles of integrins in hemostasis and immune responses are particularly important owing to their abundant expression in platelets and lymphocytes (3, 4). Abnormal expression and hyperactivity of integrins can lead to thrombotic diseases, impaired immune responses, cardiovascular diseases (CVDs), and elevated tumor metastasis (5, 6). Integrins and the associated pathways have therefore been recognized as appealing targets for therapeutic intervention and potential biomarkers for clinical diagnosis. In particular, the inside-out integrin signaling pathway that triggers integrin activation has recently emerged as an attractive target for therapeutic purposes. Integrin activation through this pathway is triggered by the association of adaptor proteins talin and kindlin to integrin on the cytoplasmic tail of its β subunit. In lymphocytes, this pathway requires a small GTPase, RAP1, and its effector <u>R</u>ap1-<u>i</u>nteracting <u>a</u>daptor <u>m</u>olecule (RIAM) (7–10). Upon activation by FAK and Src, RIAM translocates to the PM by binding to RAP1 and PIP2 (3, 11, 12). PM-anchored RIAM can recruit talin and also activates talin by disrupting its autoinhibition and dimer configuration, thus promoting talin-induced integrin activation (13–16).

Talin possesses a FERM-folded head domain of four subdomains (F0, F1, F2, and F3) and an extended rod region containing 13 helical domains (R1-R13) followed by a dimerization helix (DH), which is essential for the natural dimeric configuration of talin (17–19). RIAM mediates the conformational activation and PM translocation of talin to induce integrin activation properly. Both events require an amino-terminal helical segment of RIAM. The N-terminal, 21-residues segment of RIAM (residues 5-25), known as Talin-Binding Site (TBS) binds talin head domain in the F3 subdomain. This interaction activates talin by sterically occluding the interaction of the F3 subdomain with the autoinhibitory R9 domain (15). Interestingly, despite the close proximity of the TBS binding site and integrin binding site on the F3 subdomain, association of TBS with F3 is compatible with the F3:integrin interaction and therefore promotes integrin activation (13). Moreover, RIAM TBS also binds talin rod region in a kinked helical configuration, and this interaction is required for the translocation of talin to the PM (8, 20). This rare multi-specificity of the TBS peptide sheds light on the development of a new class of therapeutic agents that suppress integrin activity by targeting talin intracellularly using peptidomimetic inhibitors.

The major strengths of peptidomimetic drugs are good efficacy, safety, low attrition rate, and affordability (21). However, development of peptidomimetic drugs is often discouraged due to the intrinsic instability and low cell permeability (22, 23). The helical configuration of the peptide segment in the bound state may enhance its membrane permeability. However, this configuration often becomes disordered in the synthetic peptide fragment, thus reducing its membrane permeability (24, 25). To overcome the limitations of the peptidomimetic inhibitors, researchers have sought to rigidify and preorganize peptide scaffolds to stabilize the helical conformation. A molecular “staple” has been shown to improve the peptide helicity by covalently connecting two spatially-neighboring side chains in the helical configuration. Particularly, a hydrocarbon linker has been shown to significantly improve the scaffold stability and cell permeability of helical peptides (26–28). TBS adopts an α-helical conformation upon associating with both talin rod and talin head domains, making it an ideal target for stapling modification, in which a pair of sterically adjacent residues in the helical peptide can be crosslinked using a hydrocarbon linker. Moreover, because of the close proximity of the TBS binding site and integrin binding site on talin head domain, a properly designed hydrocarbon molecular staple in TBS is expected to mask the adjacent binding site of integrin on talin head and block integrin:talin interaction. In addition, a stapled TBS peptide is expected to competitively inhibit RIAM-mediated talin translocation to the PM. Thus, TBS-derived stapled peptidomimetics, as potential talin inhibitors, may achieve a high potency in inhibiting talin-induced integrin activation intracellularly in a “double-hit” manner.

Here we report the development of the first peptidomimetic talin inhibitor derived from TBS based on its unique structural, biochemical, and functional properties (20). The stapled peptide possesses a stronger affinity to talin than that of TBS and exhibits excellent cellular permeability with minimal cell toxicity. Importantly, the stapled peptide inhibits integrin:talin interaction and integrin activity inside the cells in a talin-dependent manner. We also determined the crystal structure of the stapled peptide bound to talin rod R7R8 domains. The crystal structure reveals that the stapled peptide binding site overlaps with the TBS binding site, further supporting its inhibitory effect on talin-induced integrin activation. This study provides the first example of a “double-hit” design strategy for peptidomimetic inhibitors that target two critical events in one signaling pathway. Our results offer new insight into the design of a new class of peptidomimetic inhibitors that target talin-induced integrin activation, and strongly supports the feasibility and efficacy of TBS-derived stapled peptides as new anti-integrin agents.

## Results

### The molecular staple strongly enhances TBS:talin interactions

Previous structural studies revealed that the TBS fragment binds talin head and talin rod in a helical configuration, and serve functions in activating talin and recruiting talin to the PM, respectively(13, 20). This helical configuration in the bound TBS peptide, however, is not stable in the unbound TBS peptide (20). To rigidify the helical scaffold of the TBS peptide to mimic the bound state, we designed a C8 hydrocarbon staple that connects two residues that are one helical turn apart (residues 19 and 23, **Fig. 1A**).

**Figure 1.**
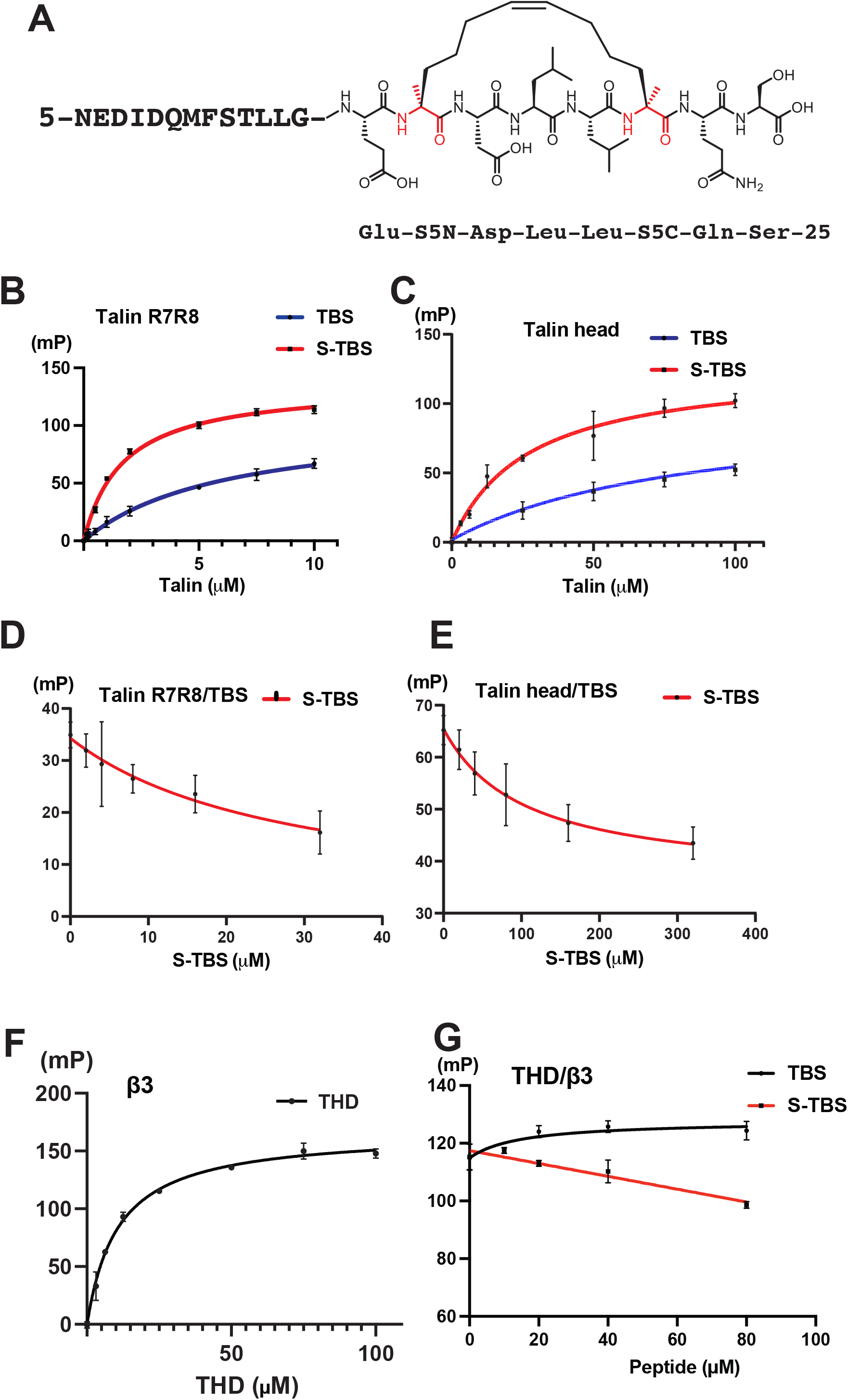
Interaction of S-TBS and talin. **A.** Structural formula of S-TBS. Residue numbers correspond to that in the RIAM sequence. **B.** Dissociation constants of TBS (red) and S-TBS (blue) with talin rod R7R8 domains were measured by FP assays using fluorophore-labeled TBS/S-TBS. **C**. Dissociation constants of TBS (red) and S-TBS (blue) with talin head domain were measured by FP assays using fluorophore-labeled TBS/S-TBS. **D**. Competitive binding of S-TBS against TBS to talin rod R7R8 domains is demonstrated by FP of fluorophore-labeled TBS. **E**. Competitive binding of S-TBS against TBS to talin head domain is demonstrated by FP. **F.** Dissociation constant of integrin β3 with talin head domain THD405 (residues 1-405) was measured by FP assays using fluorophore-labeled β3 peptide. **G**. Competitive binding of S-TBS (red) and TBS (blue) against integrin β3 to THD405 is demonstrated by FP of fluorophore-labeled β3.

To assess the impact of the C8 molecular staple on the binding affinities of TBS with talin, we measured dissociation constant (*K_d_*) of TBS or S-TBS peptides with talin head domain (residues 1-405) and with talin rod R7R8 domains using fluorophore-labeled peptides and recombinantly expressedtalin proteins by fluorescent polarization (FP) assays. TBS binds to talin head domain with a *K_d_* of 88 μM, and binds to R7R8 domains with a *K_d_* of 6.2 μM, respectively (**Fig. 1B, 1C, & Table 1**). These results are consistent with the previous studies where the mid-micromolar binding was observed with the talin head and low-micromolar binding was observed with the talin rod (13, 20). Nonetheless, affinities of S-TBS with talin head domain and R7R8 domains are significantly greater than that of TBS (3.3-fold stronger in binding to talin head and 3.6-fold stronger in binding to R7R8; **Fig. 1B, 1C, & Table 1**). We interpret that this gain of affinity is attributed to the constitutive helical configuration of S-TBS which is stabilized by the molecular staple, and the enhanced hydrophobic interaction between S-TBS and talin. The enhanced binding of S-TBS to talin further supports our design strategy of using stapled helical peptides as a potent competitive inhibitor. The competitive binding assays also indicate that the S-TBS peptide binds to both talin rod and talin head competitively against the TBS peptide (**Fig. 1D, 1E**)

**Table 1.**

| $K_d$ (μM) | | $K_d$ (μM) | 95% CI (μM) | $R^2$ |
| --- | --- | --- | --- | --- |
| Talin head domain | S-TBS | 27 | 19 - 38 | 0.9599 |
|  | TBS | 88 | 45 - 211 | 0.9537 |
| | β3 | 10 | 9.5 - 11 | 0.9958 |
| Talin rod (R7R8) | S-TBS | 1.7 | 1.5 - 2.0 | 0.9900 |
|  | TBS | 6.2 | 4.5 - 8.8 | 0.9825 |
| | β3 | N/A | N/A | N/A |

### S-TBS competes with integrin for binding talin head domain

The interaction of the TBS segment of RIAM with talin head domain activates talin by releasing it from autoinhibition. Despite their adjacent interfaces, the TBS:talin interaction is compatible with the integrin-β:talin interaction (13). To assess the impact of the molecular staple in S-TBS on integrin-β:talin interaction, we first measured the *K_d_* of FAM-labeled integrin β3 tail that includes the juxtamembrane region and the talin-binding NPLY motif (residues 720-750) with a purified talin head domain protein, THD430 (residues 1-430). We determined the *K_d_* of THD430 and β3 to be 10 μM by FP, which is more than 20-fold stronger than the previously reported F3:β3 affinity (**Fig. 1F**)(29), suggesting the FERM configuration of the full-size talin head domain stabilizes the proper folding of the F3 subdomain for integrin binding. We then assess the impact of TBS and S-TBS peptides on β3:THD interaction. No inhibitory effect was observed with the TBS peptide, whereas S-TBS exhibits a dose-dependent inhibition on talin:integrin-β3 interaction (**Fig. 1G**). These results are consistent with the compatible binding of TBS and β3 with talin head domains, and more importantly, validate our design strategy that the protruding molecular staple interferes with the binding of integrin β3 to talin head domain.

### S-TBS exhibits stable helicity and much greater membrane permeability than TBS

A common challenge in designing peptidomimetic drugs that target intracellular proteins is poor cell membrane permeability, largely due to their extended, flexible configuration and large size. Although the TBS peptide is compact and helical when bound to either talin head or rod domains, the isolated TBS fragment is rather flexible in solution (**Fig. 2A**). To examine the effect of the molecular staple on stabilizing the scaffold of the helical peptide, we assessed the helicity of S-TBS in solution using circular dichroism (CD). Indeed, S-TBS exhibits a dominant helical configuration in solution (**Fig. 2A**) consistent with our expected effect of the C8 molecular staple. We further assess the effect of the C8 staple on the cell permeability of S-TBS, we incubated fluorophore-labeled TBS or S-TBS peptides with Chinese Hamster Ovary (CHO) cells at various concentrations and examined peptide uptake by fluorescence-activated cell sorting (FACS). S-TBS treated cells exhibited significantly a stronger signal than that of the TBS peptide (**Fig. 2B**). We then performed the permeability assay at lower doses of peptides and analyzed the permeability for statistical significance. S-TBS exhibits significantly higher cell uptake at 0.5 μM. In contrast, TBS requires a concentration of at least 5 μM for cell uptake (**Fig. 2C**). We further assessed the cell viability upon treatment of the peptides. No significant toxicity in cell culture was observed in S-TBS at 40 μM concentration (**Fig. S1**). This result suggests that the hydrophobic C8 staple effectively stabilized the compact, helical configuration of the TBS peptide, and significantly improves the cell permeability of S-TBS. This data support that a properly designed hydrophobic staple in a helical peptide is able to overcome a major obstacle of poor permeability in peptidomimetic drug design.

**Figure 2.**
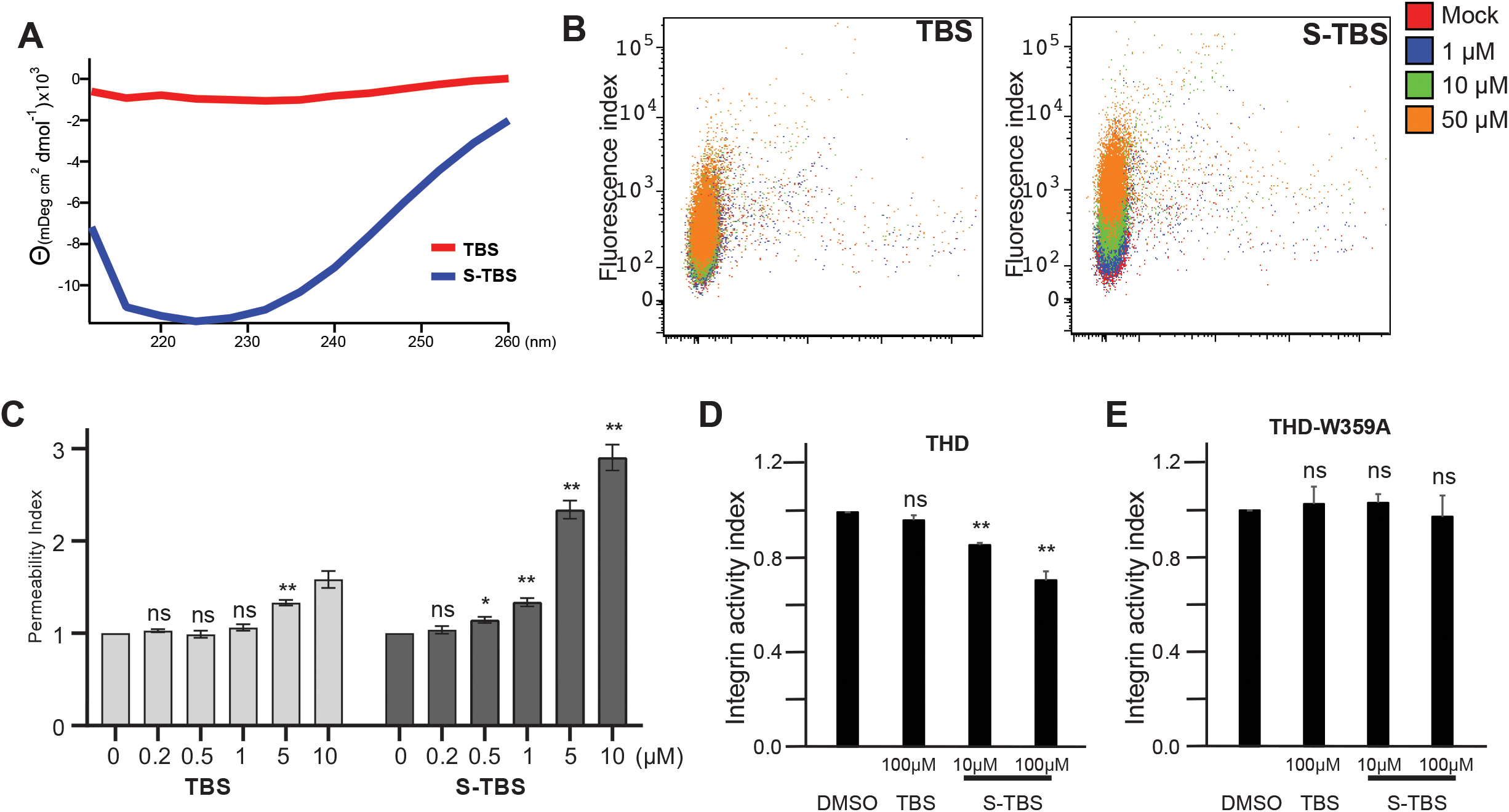
Conformational and functional characterization of S-TBS. **A.** Circular dichroism spectrometry of TBS and S-TBS in solution. **B.** Fluorescence signal of cells treated with fluorophore labeled TBS *(left)* or S-TBS *(right)* at indicated concentrations. **C.** Permeability indices of TBS *(left)* and S-TBS *(right)* measured by FACS. **D.** Integrin activity indices of αIIbβ3 in CHO-A5 cells transiently expressing wild type talin head domain (THD), and treated with TBS or S-TBS at the indicated concentrations. **E.** Integrin activity indices of αIIbβ3 in CHO-A5 cells transiently expressing talin head domain with W359A mutation (THD-W359A), and treated with TBS or S-TBS at the indicated concentrations. ns: no significance; *:*p*< 0.05; **:*p*< 0.01. Three independent experiments were performed for B, C, D and E. The data were analyzed with paired two-tailed t-test.

### S-TBS inhibits talin-induced integrin activation

We have shown that S-TBS can inhibit talin:integrin interaction *in vitro,* and processes excellent cell permeability. Later, we examined whether S-TBS is able to inhibit talin-induced integrin activation in cells. We transfected a CHO cell line that stably expresses integrin α_IIb_β_3_ (CHO-A5) with GFP-tagged talin head domain (GFP-THD) and examined the integrin activity using an active conformation-specific antibody PAC-1 by FACS assays (16, 20). S-TBS significantly suppresses activity in a dose-dependent manner, whereas the TBS peptide exhibits no inhibitory effect at 100 μM concentration (**Fig. 2D**). We further verified that the inhibitory effect of S-TBS on integrin activation is talin dependent. We examined S-TBS for inhibitory effect in CHO-A5 cells transfected with an inactive talin head mutation (THD-W359A). We observed no reduction of integrin α_IIb_β_3_ activity in cells treated with either TBS or S-TBS peptides (**Fig. 2E**), suggesting that integrin activity is insensitive to S-TBS inhibition in the absence of active talin. This result confirms that S-TBS inhibits integrin activation in a talin-specific manner. The inhibitory effect of S-TBS is also consistent with its competitive binding to talin head against integrin β3.

### Crystal structure of S-TBS in complex with talin R7R8 domains

S-TBS exhibits enhanced binding affinity with talin. To confirm that S-TBS adopts the helical conformation as seen in the talin-bound TBS peptide, we determined the crystal structure of S-TBS in complex with talin R7R8 domains (**Table 2**). The unit cell parameters and crystal packing of the S-TBS:R7R8 complex are highly similar to that of the TBS:R7R8 complex (PDB ID: 4W8P). Moreover, the 1.85-Å resolution structure reveals that S-TBS binds talin rod at the same interface of the R8 domain in a helical conformation similar to the bound TBS peptide (**Fig. 3A, 3B**). Most of the S-TBS residues, including the C8 staple connecting residues 19 and 23, are well resolved in the crystal structure (**Fig. 3C**).

**Figure 3.**
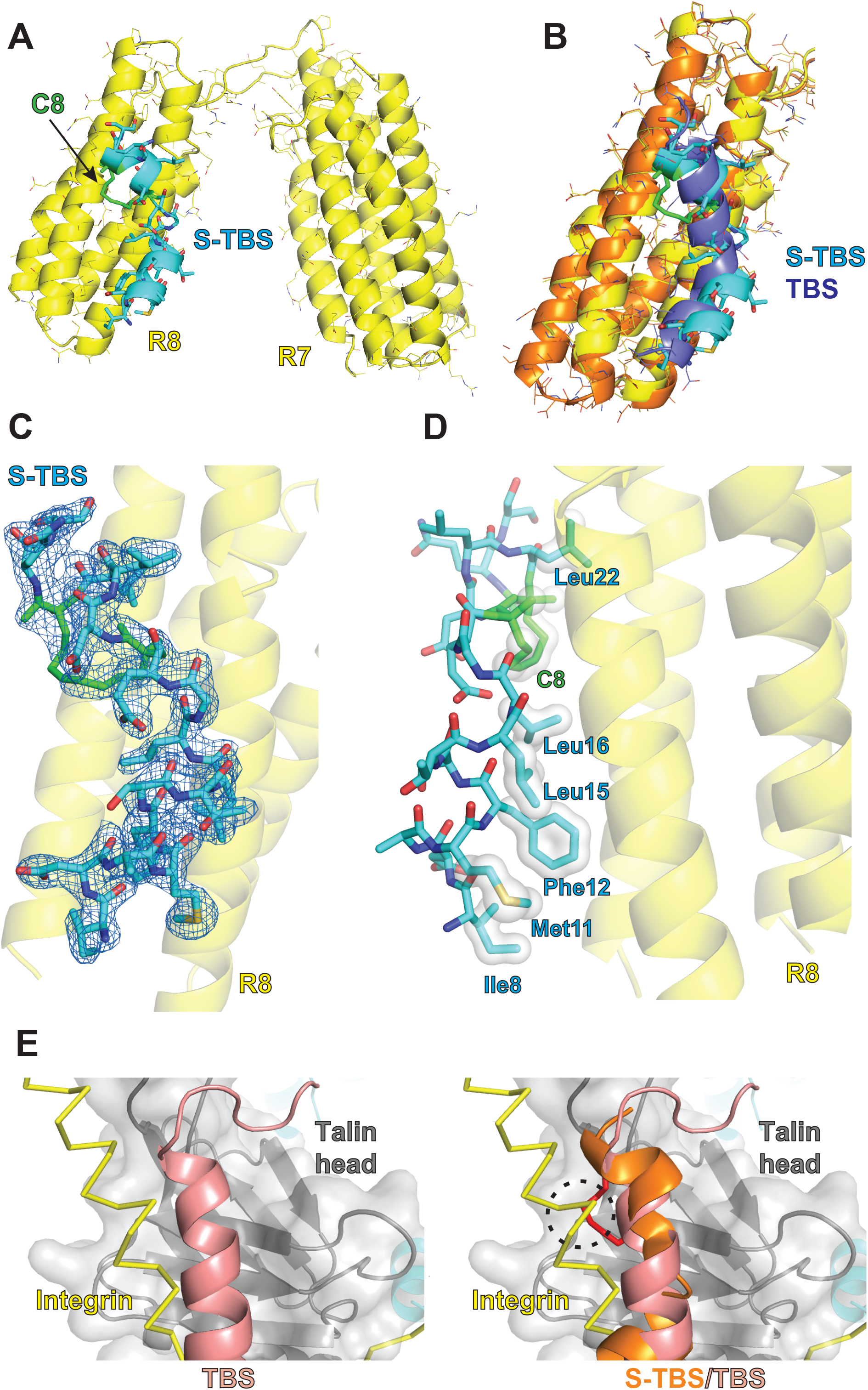
Crystal structure of S-TBS bound to talin rod. **A.** Crystal structure of S-TBS in complex with talin rod R7R8 domains. Talin domains are colored in yellow; S-TBS is colored in cyan, with the C8 linker colored in green. **B.** Superposition of the S-TBS bound R8 domain and TBS bound R8 domain (PDB: 4W8P). TBS:R8 complex is colored in orange (R8) and blue (TBS). **C.** 2Fo-Fc Electron density of the bound S-TBS contoured at 1.2σ is shown in a blue mesh. **D.** The C8 hydrocarbon linker and side chains of the residues that form the hydrophobic interface of S-TBS are shown in light gray surface representation.**E.** *Left:* Superposition of the F3 domain in RIAM-TBS:talin-F3 (PDB code: 2MWN. F3 in grey; TBS in salmon) and β1D:talin-F3 (PDB code: 3G9W. F3 in grey; integrin β1D in yellow) complexes. *Right:* Substitution of bound TBS by S-TBS peptide (orange). The collision of the C8 staple (red) and integrin β is indicated by a dashed circle.

Interestingly, although bound S-TBS also exhibits a kinked-helical conformation, the helix “turns” at residue Leu15 instead of Ser13 as seen in the bound TBS, thus allowing the side chain of Leu15 to make hydrophobic contacts with β-carbon of Thr1496 and Ala1499. This interaction compensates the loss of Glu18(TBS):Arg1510(R8) interaction in the TBS:R7R8 structure. Indeed, the S-TBS peptide contacts the R8 domain at the interface that consists of mostly hydrophobic residues, including Ile8, Met11, Phe12, Leu15, Leu16, Leu22, and the C8 staple (**Fig. 3D**). The crystal structure reveals that the binding of S-TBS to the R8 domain is dominated by hydrophobic interactions, and elucidates the inhibitory mechanism of S-TBS on RIAM:talin interaction via steric hindrance. Moreover, although structural analyses suggest that the association of juxtamembrane region of integrin and RIAM TBS with the talin head domain are compatible (13), replacement of the TBS peptide with the S-TBS peptide that adopts the similar helical conformation results in an overlap of the C8 staple and β3 (**Fig. 3E**).

## Discussion

Integrins play important roles in nearly all physiological events, and the integrin signaling pathway has become an appealing target for therapeutic intervention and a potential biomarker for clinical diagnosis. Currently, the most common anti-integrin strategy is to antagonize its extracellular ligand binding (5). Notably, all anti-integrin antagonists that have been approved for therapeutics target integrins that are abundantly expressed in blood cells (αIIbβ3, α4β1, α4β7, etc.)(30). Although these antagonistic agents show effectiveness in treating targeted diseases such as CVD and multiple sclerosis, they also cause severe adverse effects, some of which are due to paradoxical agonism of target integrins, and ligand-free integrin signaling (31–35). Therefore, alternative therapeutic approaches are in dire need to target integrin function more specifically with minimal adverse effects. Talin is required for integrin activation through its specific interaction with integrin cytoplasmic tail in the β subunit. We modified the TBS peptide with a hydrocarbon linker to stabilize the helical conformation of the peptide and enhance its binding affinities with talin. The extended hydrocarbon link in the bound, stapled peptide masks the binding site of integrin β subunit on talin head domain. The stapled TBS peptide also suppresses the interaction of RIAM and talin rod region that is required for talin translocation to the PM in leukocytes. Thus, binding of the stapled peptide to both talin head region and talin rod domain cooperatively inhibits talin-induced integrin activation (**Fig. 4A**). In addition, during integrin-mediated cell adhesion, activated integrin is linked to cytoskeleton via an integrin-talin-vinculin-actin axis, in which vinculin associates with talin via the D1 domain and with actin via the tail region (36–38). The TBS peptide is also capable of interacting with the D1 domain of vinculin, thus competing against talin for binding to vinculin (38). Therefore, this peptide may impact three essential events in the integrin signaling pathway including talin translocation to the PM, talin activation, and talin-vinculin interaction. The unique multi-specificity of TBS poses a unique opportunity for developing a therapeutic agent that may suppress integrin activity intracellularly and avoid agonism.

**Figure 4.**
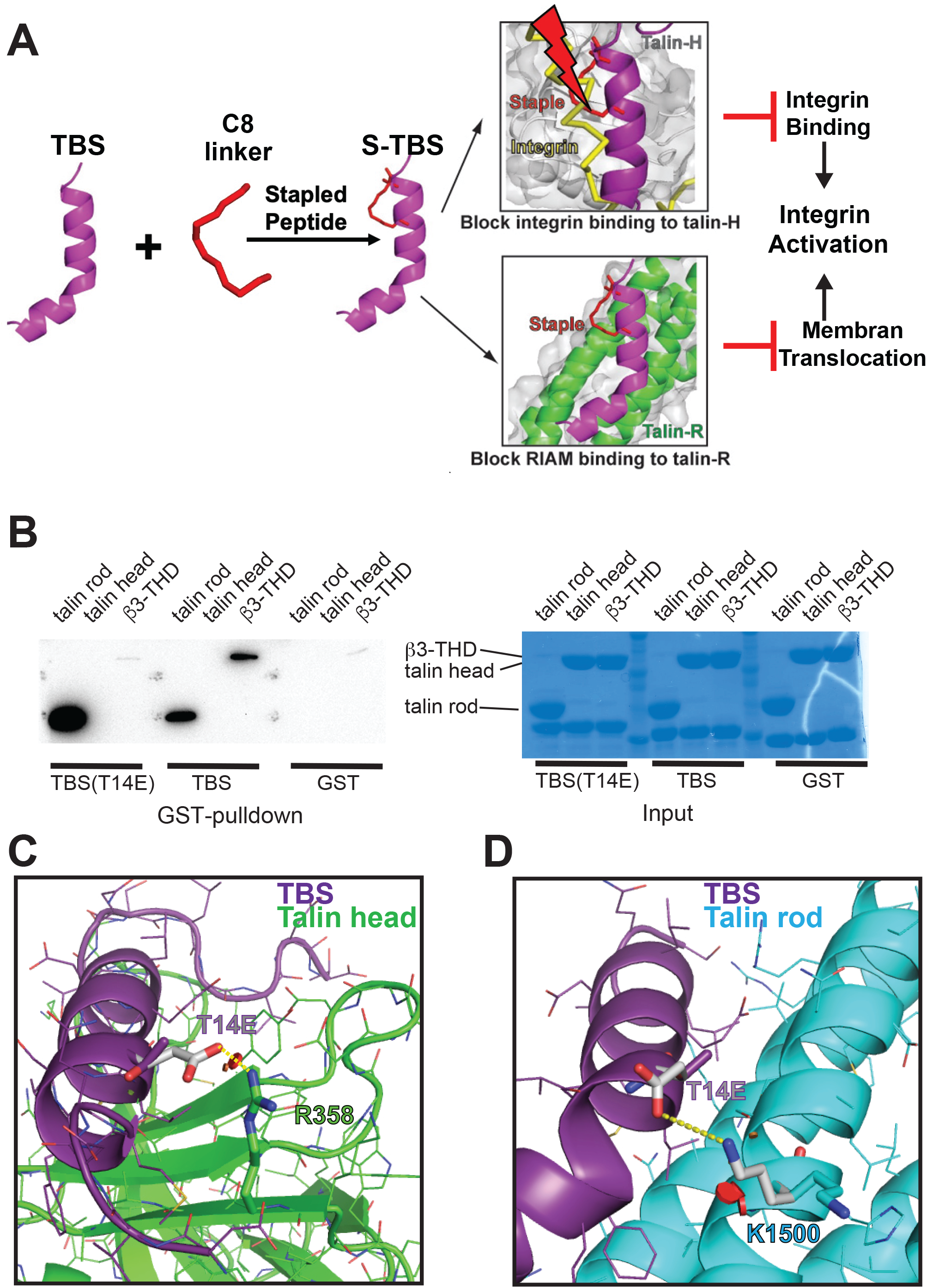
Inhibiting talin-induced integrin activation by stapled TBS peptide. **A.** TBS peptide (purple) is modified by a hydrocarbon (red) link to stabilize its helicity. The enhanced affinity of the stapled TBS peptide occludes RIAM from binding to both talin head domain (gray) and talin rod region (green), and masks the binding site of integrin β subunit (yellow), cooperatively inhibiting talin-induced integrin activation. **B.** GST-pulldown of talin rod (R7R8 domains), talin head (residues 1-430), or β3-THD (integrin β3-talin head domain fusion protein) by GST-TBS or GST-TBS(T14E). **C**. T14E (gray) in TBS is predicted to form a salt bridge interaction with Arg358 in talin head, **D**. T14E (grey) in TBS is also predicted to form a salt bridge interaction with Lys1500 of talin R8 domain (cyan) in a different rotamer (gray).

Our data demonstrated that, although the inhibitory efficacy of S-TBS ontalin:integrin interaction appears to be moderate, its effect is much more significant on talin-mediated integrin activation. We examined the talin:integrin interaction using a fluorophore labeled integrin β3 fragment that includes both the juxtamembrane (JM) region and the downstream talin-binding NPLY motif. Previous structural studies have indicated that both JM and the NPLY motif interact with talin head domain (29, 39). In the FP assays, although S-TBS can outcompete the JM region for talin binding, the flexibility of theβ3 fragment may still allow the NPLY motif to bind talin with a reduced affinity. Nevertheless, talin-mediated integrin activation requires not only the interaction of the NPLY motif and talin head domain, but also a proper association of the β-JM region and talin head domain, and more importantly, the catalytic interaction of the talin F1 loop with the JM regions of α- and β-subunits of integrin (19). Thus, the interaction of JM and talin head is also essential for integrin activation as it properly positions the JM region near the talin F1 loop. Therefore, the apparent inhibitory effect of S-TBS can be better demonstrated by the cell-based integrin activity.

Nevertheless, in order to optimize the efficacy, gain-of-function (GOF) mutations may be introduced in TBS to enhance its interaction with talin. It has been speculated that TBS may synergize the integrin-β:talin interaction. Interestingly, our data indicate that a chimeric protein that fuses an integrin β3 segment to talin head domain (β3-THD) interacts with GST-TBS much more strongly than THD alone (**Fig. 4B**). Indeed, synthesized TBS peptide also enhances the interaction of THD and FAM-labeled β3 peptide in the FP assay (**Fig. 1G**). These results support the synergistic interaction of integrin-β and talin head by TBS. Furthermore, our structural analyses suggest that a T14E mutation in TBS is expected to enhance TBS interaction with both talin head and rod domains. Substitution of Thr14 by a glutamate residue may generate a new salt-bridge interaction with talin head domain through Arg358 (**Fig. 4C**). It is worth noting that Arg358 is essential for the interaction of talin and integrin(40), and TBS-T14E is expected to compete against integrin for binding to talin as a result. Moreover, the T14E mutation is also predicted to generate a new salt-bridge interaction with talin R8 domain through Lys1500 (**Fig. 4D**). Indeed, the T14E mutation significantly diminishes the interaction of TBS with β3-THD (**Fig. 4B**), and exhibits stronger interaction with talin R7R8 domains (**Fig. 4B**). Thus, T14E, and other potential GOF mutations may be introduced into the “second generation” of experimental stapled peptides. The efficacy of the peptidomimetic inhibitor may also be further improved by optimizing the linker size, chemical composition, and connecting sites of the molecular staple. Nevertheless, further investigations are required to assess drug stability, facilitate tissue specific drug delivery, and resolve other issues related to drug development.

In summary, based on the unique functional properties of TBS, we designed the TBS-derived, stapled peptidomimetic compound as a talin inhibitor that targets talin-mediated integrin activation. We developed S-TBS as a “proof-of-concept”, an experimental agent that validates the “double-hit” design strategy which, to our knowledge, is the first structure-based designed inhibitor of talin. S-TBS exhibits excellent cell permeability without significant cell toxicity. Importantly, S-TBS is capable of inhibiting integrin activity by specifically targeting talin intracellularly. We anticipate that this approach of inhibitor design by targeting multiple events in a pathway would not only lead to the discovery of novel anti-integrin drugs but also help in identifying new intracellular targets and facilitate the development of the corresponding peptidomimetic inhibitor to improve the therapeutic outcomes.

## Materials and Methods

### Plasmid construction, protein purification, and GST pull-down

Talin R7R8, talin head domain, β3-THD chimera, and GST-TBS were expressed and purified as described previously (18, 20). Point mutations were constructed using a site-directed mutagenesis method. For *in vitro* pull-down assays, purified GST-TBS, wild type or T14E, were immobilized on glutathione agarose beads and then incubated with purified His-tagged talin proteins in reaction buffer (20 mM Tris-HCl pH 7.5, 100 mM NaCl, and 2 mM DTT) to a total volume of 250 μl on a rotator for 1 hr at 4°C. The bound proteins were washed three times in 500 μl of the lysis buffer and were eluted using an elution buffer (20 mM Tris-HCl pH 7.5, 100 mM NaCl, 2 mM DTT, and 10 mM reduced glutathione) at 4°C. The proteins were resolved by SDS-PAGE and detected by Coomassie staining or Western blotting. The Immobilon-P transfer membranes (EMD Millipore) were blocked with TBST buffer (20 mM Tris-HCl pH 7.5, 150 mM NaCl, 0.1% (v/v) Tween 20) containing 5% (w/v) BSA for 1 hour and then incubated with anti-His (Sigma) or anti-GFP antibody (Clontech) for 1 hour at room temperature followed by a second incubation with HRP-conjugated secondary antibody (Santa Cruz Biotechnology). The blots were visualized with the SuperSignal West Pico Chemiluminescent Substrate (Thermo Scientific) and detected using the FluorChem E imager (ProteinSimple).

### Crystallization and structure determination method

The talin rod domains R7R8 was purified as described in the previous study (20). The S-TBS peptide (New England Peptide), possessing mouse RIAM residues 5-25 with a C8 linker between residues 19 and 23, was solubilized in 20 mM Tris pH8.0, 100 mM NaCl with 50% DMSO. Talin R7R8 (35 mg/mL) was incubated with the S-TBS on ice at a 1:3 molar ratio prior to the crystallization setup. S-TBS:R7R8 complex was crystallized using the hanging-drop vapor diffusion method at room temperature in 100 mM NaCl, 20% (w/v) polyethylene glycol 3350 and 20% (v/v) ethylene glycol. X-ray diffraction data were collected at the APS NECAT 24-ID-E beamline. The collected datasets were indexed, integrated and scaled with XDS (41). Data collection and refinement statistics are listed in **Table 2**. The crystal structure was determined by molecular replacement using R7R8 structure (PDB: 4w8p) as a search model, the model was iteratively built in COOT(42) and refined in PHENIX and REFMAC (43). To build the model of the stapled peptide, Met19 and Thr23 were replaced by irregular amino acid MK8 and the side chain was linked. The final atomic coordinates and structure factors have been deposited to Protein Data Bank with accession number 7V1A.

**Table 2:** Crystallographic data collection and refinement statistics.

|  |  |
| --- | --- |
| **Data collection** |  |
| Light Source | NSLS-II AMX |
| Wavelength (Å) | 0.97918 |
| Space group | P2<sub>1</sub>2<sub>1</sub>2 |
| Cell dimensions |  |
| *a*, *b*, *c* (Å) | 60.86, 104.33, 48.23 |
| $\alpha$, $\beta$, $\gamma$ (°) | 90, 90, 90 |
| Resolution (Å) | 104.33~1.85 (1.88~1.85) |
| *R*<sub>merge</sub> (%) | 3.3 (154.6) |
| *I*/$\sigma$ | 22.6 (1.0) |
| CC(1/2) | 99.9 (47.5) |
| Completeness (%) | 99.4 (91.9) |
| No. reflections | 27043 (1512) |
| Multiplicity | 6.5 (6.1) |
| **Refinement** |  |
| Resolution (Å) | 52.57~1.85 |
| *R*<sub>work</sub>/*R*<sub>free</sub> (%) | 21.54/25.38 |
| No. atoms |  |
| Protein | 2172 |
| Ligand/ion | 152 |
| Water | 144 |
| Wilson B factor (Å<sup>2</sup>) |  |
| Protein | 60.6 |
| Peptide | 87.44 |
| R.M.S deviations |  |
| Bond lengths (Å) | 0.010 |
| Bond angles (°) | 1.343 |

### Fluorescent polarization

We mixed purified talin proteins at a series concentration (R7R8: 0.2 μM to 10 μM; THD/β3-THD: 50 μM to 100 μM), or buffer only, with 20 nM FITC-labeled S-TBS (New England Peptide) in a buffer containing 20 mM Tris, pH 8.0, 100 mM NaCl, and 2 mM DTT. After 5 min of incubation, we added 20 μl of the reaction mix to individual wells of a 384-well assay plate (Corning) and then measured fluorescence polarization at room temperature using a plate reader (Perkin Elmer Envision Plate Reader). We used FITC FP 480 filter as an excitation filter, and a pair of FITC FP P-pol 535 filters as emission polarization filters. FP signals were normalized against the buffer-only background and then fit to a single-site (saturating) binding model using Prism 9. Competitive FP assay was performed by titrating a series concentration of unlabeled S-TBS into a solution mixture containing 10 μM R7R8 (or 40 μM THD) and 20nM FAM-TBS. Competitive binding of S-TBS to R7R8 and THD against FAM-TBS are indicated by the reduction of FP signals.

### Cell permeability assay

CHO-A5 cells were plated in 6-well dishes and then treated with fluorophore-labeled TBS or S-TBS peptides at indicated concentrations for 30 min at 37°C. The cells were then washed twice with phosphate-buffered saline (PBS), detached by adding trypsin-EDTA and harvested by spinning at 2,000 rpm for 5 min. The cells were resuspended in PBS and quantified with an LSR flow cytometer (525/50 filter, 488 nm laser) using 30,000 cells per measurement.

### Integrin activation assays

CHO-A5 cells were transfected with talin-head domain, WT or W359A, and incubated for 24 h at 37°C.

The cells were split to achieve equal expression level of talin head, and subsequently treated with TBS peptide or S-TBS for 30 min at 37°C and washed twice with PBS buffer. The cells were then detached by adding trypsin-EDTA, and harvested by spinning at 2,000 rpm for 5 min. The cells were incubated in Tyrode’s buffer (136.9 mM NaCl, 10 mM HEPES, 5.5 mM Glucose, 11.9 mM NaHCO_3_, 2.7 mM KCl, 0.5 mM CaCl_2_, 1.5 mM MgCl_2_, 0.4 mM NaH_2_PO_4_, pH 7.4) containing PAC1 antibody (BD) for 1 h at 30 °C and then stained with Alexa 647 goat anti-mouse IgM (Invitrogen) for 30 min at RT. The cells were quantified with an LSR flow cytometer (660/20 filter, 640 nm laser) using 30,000 cells per measurement.

### Cell viability assays

HEK-293, MCF-7, or HeLa cells were plated in 24-well plates with proper coatings at 20% confluency. Cells were treated with TBS or S-TBS peptides, or Dasatinib, at the indicated concentration. Cell viability will be determined after 24 hours, 48 hours, and 72 hours by using the Deep Blue Cell Viability^TM^ kit (Biolegend).

### Data Availability

The data that support the findings of this study are available from the authors on reasonable request, see author contributions for specific data sets.

### Accession Numbers

The accession number for the atomic coordinates and structure factors reported in this paper is PDB: 7V1A (R7R8:S-TBS)

## Author Contributions

T.G. purified proteins used in the FP assay. T.G. and E-A.C. performed FP assay. E-A.C, T.G. and P.Z. performed the cell permeability assay. P.Z. and T.G. performed the protein production and crystallization. P.Z.and J.W. processed the crystallographic data and determined the structures. T.G and P.Z. performed pull-down assay. E-A.C performed the cell-based functional experiments. T.G., E-A.C, and P.Z. contributed to the manuscript preparation. J.W. supervised the project and was the principal manuscript author.

## Acknowledgments

We thank Leland Mayne of The Biophysical and Structural Biology Core at Perelman School of Medicine, University of Pennsylvania, and the beamline staff of NECAT 24-ID-E at the Advanced Photon Source, Argonne National Laboratory for technical help. This work was supported by an NIH Grant GM119560, a Pennsylvania Department of Health Grant 4100085739, and ACS RSG-15-167-01-DMC. T.G. was also supported by the Elizabeth Knight Patterson Postdoctoral Fellowship. E.A.C. was also supported by grant 5T32-CA-009035-42 to FCCC.

---

**Figure S1.**
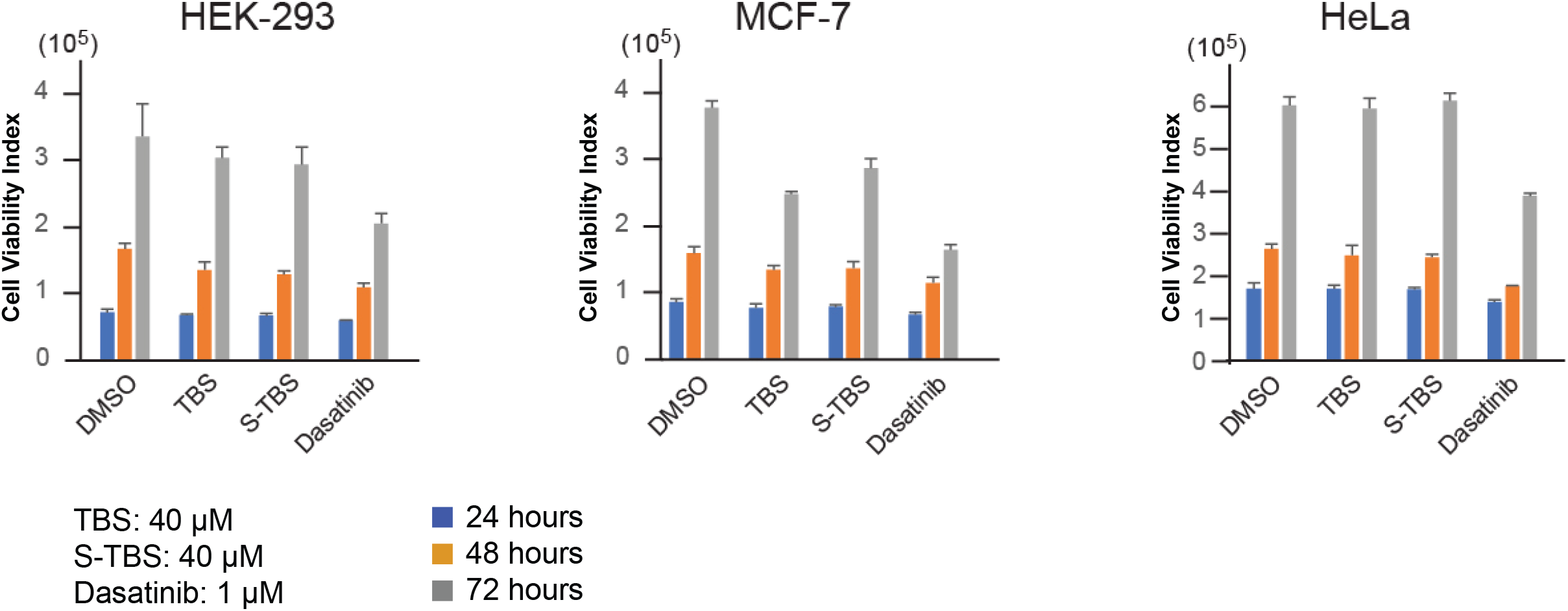
Cell viability test of TBS and S-TBS peptides. HEK-293, MCF-7, or HeLa cells were treated with 40 μM TBS or S-TBS, or 1 μM Dasatinib. Cell viabilities of each cell types were measured at 24 hrs (blue), 48 hrs (orange), and 72 hrs (gray).

